# Zooming into the genomic vicinity of the major locus for vicine and convicine in faba bean (*Vicia faba* L.)

**DOI:** 10.1101/2021.02.19.431996

**Authors:** Rebecca Tacke, Wolfgang Ecke, Michael Höfer, Olaf Sass, Wolfgang Link

## Abstract

The versatility of faba bean (*Vicia faba* L.) seed as valuable protein feed is hampered by its relatively high level of the compounds vicine and convicine (VC), which are antinutritive factors in poultry and further non-ruminant feed. The objective here was to develop the first-ever genetically low-VC winter faba bean. Hence, the low VC allele vc-, that should be the basis of a known, major locus for VC, need verification and molecular identification and be based on appropriately developed DNA-markers; the low VC feature awaited its transfer into the high-performing winter faba bean germplams. Based on bi-parental F_2_-families and isogenic lines, we thus developed highly useful SNP markers exploiting transcriptomic data. Furthermore, we fine-mapped and, based on synteny to *Medicago truncatula* and *Cicer arietinum*, we identified a candidate gene for the VC locus. A novel, genetically low VC winter faba bean population was bred. The path is now well-prepared for further marker-based breeding progress.

## Introduction

Faba bean (*Vicia faba* L.; *V.f.*) is a large-seeded, annual grain legume which is grown for food and feed. It is appreciated for its high protein content of approximately 30% (Crépon et al. 2010; Link et al. 1994), as a regional vegetal protein source and as an alternative crop to improve soil fertility and break rotation circles of pests and diseases (Köpke and Nemecek 2010; Kulak et al. 2013). Genomic tools are still underdeveloped for faba bean (Annicchiarico et al. 2017); a sequence of the huge genome of faba bean (13 Gb) is not yet available. Therefore, the high syntenic correspondence of *Vicia faba* to the sequenced genomes of *Medicago truncatula (M.t.), Cicer arietinum (C.a.)* and further legumes of the subfamily *Faboideaei* is of high importance for genomic analyses. Faba beans are cultivated in Germany on only approximately 50.000 ha (Statistisches Bundesamt 2020). One limiting factor is the presence of the antinutritive seed compounds <u>vicine and convicine</u> (VC), two pyrimidine glycosides which occur in faba bean seed with approx. 0.3% up to 1.5% in dry matter. This quantita-tive variation has a clear genetic component (Duc 1997; Frauen et al. 1984; Khamassi et al. 2013). Vicine and convicine occur in an approximate 2:1 ratio in faba bean seeds (Goyoaga et al. 2008). Faba beans contain VC in all parts of the plant (Goyoaga et al. 2008). These compounds are nearly unique to the genus *Vicia*; *Momordica charantia* (*M.c.*) is the only species outside this genus containing vicine (Gauttam and Kalia 2013; Khazaei et al. 2019).

More than 400 million people will, upon ingestion of vicine and convicine, suffer from a hemolytic anemia, called favism, caused by a human X-chromosomal inherited genetic deficiency of glucose-6-phosphate-dehygrogenase (G6PD). This human genetic condition has a delicate epidemiological connection to malaria (Arese 2006; Arese et al. 2012; Luzzatto and Arese 2018).

Vicine and convicine can also have a negative effect e.g. on laying hen and broiler performances (Guillaume and Bellec 1977; Halle 2006; Larbier and Leclercq, 1994; Marquardt et al. 1981; Munduuli et al. 1981; Muduuli et al. 1982; Naber et al. 1988; Olaboro et al. 1981a; Olaboro et al. 1981b).

Duc et al. (1989) reported the presumably monogenetically inherited <u>low vicine and convicine</u> (LVC) content of the genebank accession 1268(4)(1) from Radzikov (Poland), which has since been used in breeding and research. This accession showed a seed content of 0,046% of VC, this is about 1/10 to 1/20 of the wild type seed content. That trait is controlled by one locus, designated as VC locus, with <u>alleles VC+</u> (wild type) and <u>vc-</u> (from that genebank accession 1268(4)(1)). The low VC level in homozygous vc-/vc- faba bean genotypes prevents favism in G6PD-deficient humans (Gallo et al. 2018) and prevents dietary disadvantages if such faba beans are used as compounds of feed (Crépon et al. 2010). Heterozygosity at the VC locus causes approximately intermediate VC values.

Based on scattered hints in classical literature, vicine and convicine are assumed to be formed in the faba bean seed coat and transported into the embryo (Duc et al. 1989; Brown and Roberts 1972).

The biosynthetic pathway to vicine and convicine was unknown until recently, when Björnsdotter et al. (bioRxiv preprint, 2020) presented new, pioneering findings, which provided convincing experimental evidence that a bi-functional RIBA1 protein, which catalyzes the first step of the riboflavin biosynthetic pathway, also catalyzes the key step of the so far unresolved vicine and convicine pathway. Starting with the GTP cyclohydrolase II function of this protein, the two pyrimidine glucosides are synthesized from GTP as demonstrated by feeding isotopically labelled GTP precursor into roots of *V. faba*. However, the additional enzymatic steps, which finally lead to the compounds vicine and convicine, remain to be elucidated, although the authors propose that VC are synthesized in three additional steps from the first two intermediates in the riboflavin pathway, respectively, the first of which is probably catalyzed by an N-glycosidase described by Frelin et al. (2015). Björnsdotter et al. (2020) could also show that there is a copy of the gene encoding the RIBA1 protein at the VC locus, whose GTP cyclohydrolase II function is destroyed by a frameshift mutation in the vc- allele. This gene was consequently named VC1.

A major objective of research of VC characteristics is to facilitate breeding of faba beans which are genetically low in vicine and convicine seed content. The black and white hilum colours of faba bean seeds, a monogenic trait (Sirks 1931), can be used as morphological marker to select for LVC status. The accession 1268(4)(1), source of the LVC feature, shows white seed hilum. The linkage between the VC locus and the hilum colour locus is relatively tight (5-10cM; Duc et al. 2004; Khazaei et al. 2017). Yet, white versus black hilum is only visible late in the life of plants, when their seeds are mature. In an early attempt to overcome this limitation, Gutierrez et al. (2006) developed two CAPS markers by bulked segregant analysis, which were linked to the VC locus. Later, Khazaei et al. (2015) were able to map the VC locus as a major QTL on the large first chromosome of faba bean. The QTL mapped in an interval of 3.6 cM, flanked by two pairs of cosegregating SNP markers. Unfortunately, the two markers closest to this mapped QTL (0.8 cM) proved to be not diagnostic for the trait in a larger set of diverse genotypes.

The genetic map used for the mapping of the VC locus had been constructed with SNP markers (Khazaei et al. 2014) that were a subset of a larger set of 687 SNP markers which were used to construct a comprehensive consensus map for faba bean (Webb et al. 2016). The majority of these SNP markers had been developed based on selected mRNAseq contigs of faba bean that could be unambiguously assigned to a specific gene of *M.t.* by sequence comparisons. These markers therefore allow comparative mapping between faba bean and *M. truncatula.* Khazaei et al. (2015) used this approach to identify a region on chromosome 2 of *M. t.* with strong collinearity to the region in faba bean carrying the VC locus. A total of 340 genes were found in *M. t.* in the interval delimited by the markers flanking the VC locus. However, due to lacking knowledge about the biosynthetic pathway for vicine and convicine, it was not possible to identifiy a candidate gene for the VC locus. In an early approach to identify candiate genes for the VC locus, Ray et al. (2015) identified six RNAseq contigs that were differentially expressed between high and low VC genotypes. One of these contigs, contig 4518, was later located by Khazaei et al. (2017) in *M. t.* in the target region for the VC gene on chromosome 2. The contig 4518 corresponds to the *M. t.* gene Medtr2g009270, which is annotated as 3,4-dihydroxy-2-butanone 4-phosphate synthase. In soybean and chickpea, the same gene is annotated as RIBA 1, a bifunctional riboflavin biosynthesis protein (Khazaei et al. 2017) that has two catalytic domains, one with a 3,4-dihydroxy-2-butanone 4-phosphate synthase activity and a second one with a GTP cyclohydrolase activity. Based on contig 4518, Khazaei et al. (2017) developed an SNP marker whose segregation completely matched the segregation of VC content in a RIL population from a cross between a high and low VC genotype and in a set of 51 diverse genotypes. A comprehensive and detailed review on the current classical and molecular genetic background of the VC feature of faba bean was very recently presented by Khazaei et al. (2019).

Here, the main objectives were to demonstrate and facilitate the breeding of faba beans with low VC content. Therefore, we developed new SNP markers which were closely linked to the VC locus via fine-mapping and, if possible, locate and identify the VC gene.

Finally, we aimed to employ all our findings to develop a novel, LVC winter faba bean germplam.

## Material and Methods

### Plant material for genetic analyses

We used two pairs of <u>near-isogenic lines</u> (NILs) for mRNA analysis and as crossing parents to develop two F_2_ mapping populations (Cross1, Cross2). These two NIL pairs were themselves generated from two crosses of a <u>high vicine and convicine</u> (HVC) and a LVC parental inbred line, respectively. The first NIL pair originated from the cross between the inbred line Mélodie/2 and ILB938/2. Mélodie/2 was bred via single-seed descent (SSD) from the French LVC cultivar Mélodie, whereas ILB938/2 was bred via SSD from the HVC gene bank accession ILB938 (Khamassi et al. 2013; Khazaei et al. 2015). The second NIL pair was accordingly generated from a cross facilitated by NPZ (Norddeutsche Pflanzenzucht Hans-Georg Lembke KG). This cross included the cultivar Fabelle as the LVC parent. From both crosses, recombinant inbred lines (RILs) were bred until generation F5 (Khazaei et al. 2015). Here, one rare F5-individual per cross was identified which gave F6 offspring that still segregated for VC and included a homozygous HVC and homozygous LVC individual. These individuals were maintained by self-fertilization and became one pair of NILs. Hence, one such pair of NILs was derived from each of the two initial crosses. From Mendelian expectation, the two lines of each NIL pair should be isogenic at about 15/16 of their genome.

These were the near-isogenic parents for Cross 1:

- “Mél/2*ILB938/2-139-1 (LVC)”
- “Mél/2*ILB938/2-139-2 (HVC)”

These were the near-isogenic parents for Cross 2:

- “NPZ-848-3 (LVC)”
- “NPZ-848-4 (HVC)”

### Creation of mapping populations

To create the two F2-mapping populations, the F_1_ of Cross 1 and of Cross 2 were self-fertilized under insect-proof conditions. These two mapping populations contained N=751 (Cross 1) and N=899 (Cross 2) F_2_-individuals. The two F_2_ populations where subsequently used to generate <u>linkage map fragments</u> and to conduct fine-mapping analyses while seed coat material from their parental NIL pairs was further used for mRNA analyses.

### Plant material for breeding novel, winterhardy, LVC faba bean lines

This breeding program started in 2006 by crossing Hiverna/2 and Mélodie/2. Hiverna/2 had been bred via SSD from Hiverna, an old, very winterhardy German winter faba bean cultivar with HVC phenotype (Bundessortenamt 2019).

Until 2015 and generation BC_2_F_2_, the selection was carried out via hilum colour among F_2_, BC_1_F_2_ and BC_2_F_2_ individuals. For actual backcrossing, corresponding F_3_, BC_1_F_3_ and BC_2_F_3_ individuals were used. Starting in BC_3_F_2_, selection was based on high liquid pressure chromatography (HPLC) analyses (service provided by National Institute of Agricultural Botany in England, NIAB 2021). Further on, we employed marker assisted selection for the next backcrossing program.

### mRNA sampling, RNA-Isolation, transcriptome, and SNP analysis

For mRNA analyses, we collected seed coats of immature seeds (Figure 1), in their developmental stages four, five and six (Borisjuk et al. 1995).

**Figure 1.**
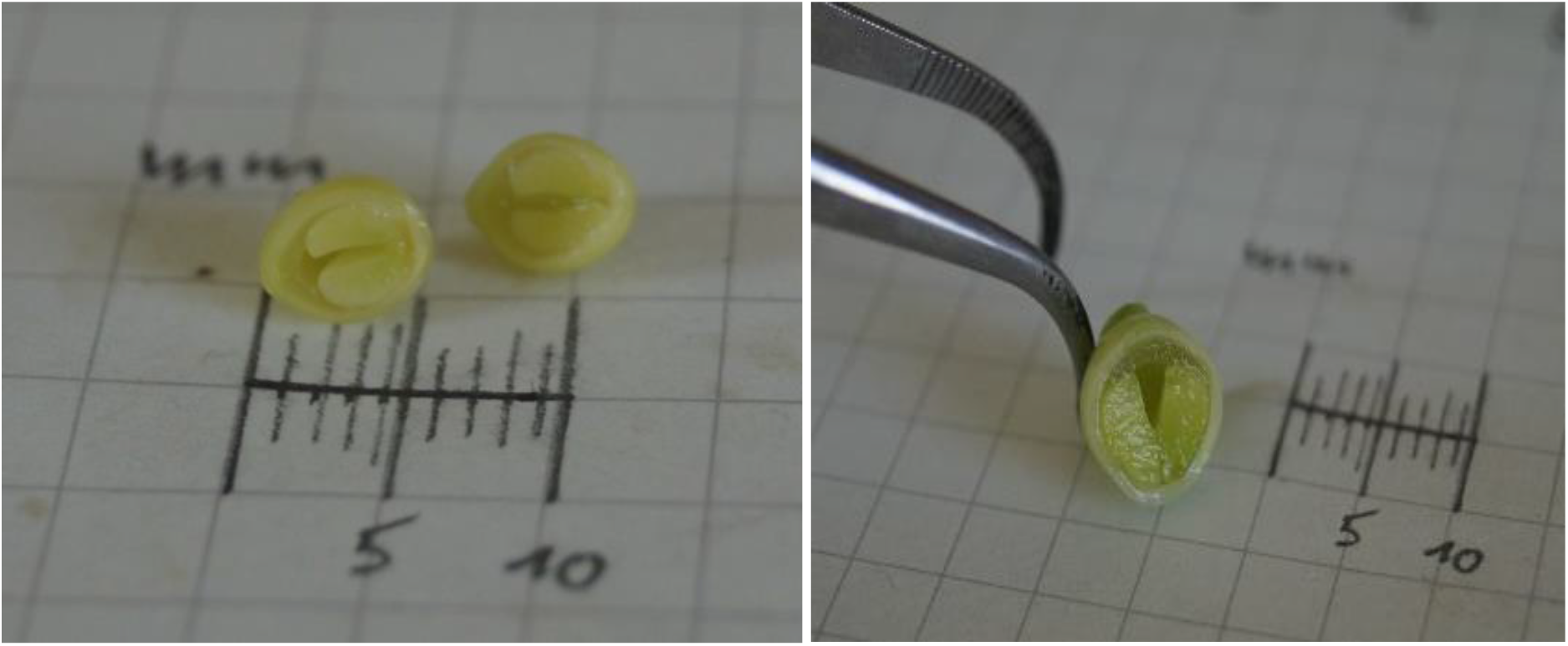
Cross-section of a *Vicia faba* seed in developmental stage 4 (left) and stage 5 (right).

The frozen tissue was ground to a fine powder under liquid nitrogen conditions using a mixer mill (MM200, Retsch, Haan, settings 30 Hz for 2 min). Total RNA was isolated with the InviTrap^®^ Spin Plant RNA Mini Kit (STRATEC SE, Birkenfeld). Aliquots of this RNA were treated with Baseline Zero™ DNase (Biozyme Scientific GmbH, Hessisch Oldendorf) according to the manufacturer’s instructions to eliminate contaminating gDNA, purified using the MinElute Cleanup Kit (STRATEC SE, Birkenfeld) and finally photometrically measured (ND-1000, NanoDrop Technologies, Wilmigton, USA). The RNA integrity number values of all RNA samples to be sequenced were determined using a 2100 bioanalyzer (Agilent Technologies, Waldbronn, Germany). Individual RNA samples for transcriptome analysis were converted into 3′-prime specific MACE libraries (Bojahr et al. 2016) and sequenced with 1x 75 b reads on a HiSeq 2000 machine (Illumina) resulting in 3,1 - 5 million quality filtered reads. For each line of the NIL pairs, RNA samples which derived from seeds of the developmental stages 4 and 5 were pooled on a ratio of 1:1 and subsequently converted into RNASeq libraries and paired-end sequenced at 2×75 b on a Hiseq2000 device yielding 64-77 million quality filtered reads per sample. Combined high-quality reads from all samples were used for a *de novo* assembly and annotation of a duplicate free testa-specific reference transcriptome as described in Santos et al. (2018). The software package JointSNVMix was used to detect single nucleotide variants (Roth et al. 2012). For SNP candidates, a locus must have had at least five reads in both samples where one of two possible bases might occur (either allele A or B). A maximum of one percent false allele reads was tolerated. NGS template preparation, sequencing and bioinformatics took place at GenXPro GmbH (Frankfurt am Main). Transcriptome data evaluation for the identification of SNPs that correlated with the VC phenotype was performed stepwise: for the RNASeq and MACE libraries separately and for each of the two NIL pairs (parents of Cross 1 and Cross 2). For each genotype, MACE-data sets derived from developmental stages 4, 5, and 6 were pooled to increase sensitivity for SNP-detection. Subsequently, those SNP candidates which were common to both NIL pairs were identified.

### SNP marker development

KASP marker development, DNA extraction and KASP analysis were conducted at TraitGenetics (Gatersleben, Germany). KASP assay designs were based on app. 50 bps to the left and right side of SNPs found via the mRNA analysis.

### Marker analysis and map construction

Throughout, the genetic linkage maps were constructed using R (R Development Core Team 2011; R version 3.5.3) and the R package ASMap (Taylor and Butler 2017). For all linkage maps, the Kosambi function was used to calculate the map distances in cM (Kosambi 1943). The functions of the package were used as described by Taylor and Butler (2017).

Missing marker data and marker segregation ratios were identified with the R Package ASMap. Markers that were monomorphic in a cross and markers showing strongly skewed segregations (−log(P) of 2.0 or higher) were excluded from further analyses. The selected SNP markers were used for the linkage map fragment construction.

An <u>initial marker set</u>, which contains markers from Webb et al. 2016, Song et al. 2017, and the group of Donal O`Sullivan, yielded the so-called <u>draft maps</u>. These were used to define the putative region of the VC locus for future analyses, especially for the transcription analysis.

The set of newly developed markers from our transcription analyses and certain markers from the initial marker set (termed <u>new marker set</u>) yielded the <u>final linkage map fragments</u>, which were used for the fine-mapping.

### Phenotyping of VC content for fine-mapping and breeding

Selected F_2_ individuals from Cross 1 and Cross 2 as well as the backcross generation BC_3_F_2_ were phenotyped (HPLC; via NIAB 2021). As checks and benchmarks for the HPLC results, F_2_ individuals with marker-deduced homozygous and heterozygous VC genotypes were identified (see Table 1).

**Table 1.** VC contents of homozygous and heterozygous F_2_ individuals of Cross 1 and Cross 2.

| Genotype of F<sub>2</sub> individuals |  | No. of individuals | VC range of HPLC values | Mean | Standard error | VC content category |
| --- | --- | --- | --- | --- | --- | --- |
| **Cross 1** | Homozygous HVC | 6 | 0.54 – 0.89 | 0.6428 | 0.0502 | HVC |
|  | Heterozygous | 8 | 0.47 – 0.49 | 0.4798 | 0.0088 | intermediate |
|  | Homozygous LVC | 5 | 0.04 – 0.05 | 0.0421 | 0.0015 | LVC |
| **Cross 2** | Homozygous HVC | 4 | 0.51 – 0.61 | 0.5520 | 0.0243 | HVC |
|  | Heterozygous | 7 | 0.32 – 0.34 | 0.3311 | 0.0080 | intermediate |
|  | Homozygous LVC | 5 | 0.02 – 0.03 | 0.0270 | 0.0020 | LVC |

### Fine mapping

For the manual fine-mapping of the sought-for VC-locus, phenotypic VC results were displayed along with the SNP genotypes of the F_2_ individuals of Cross 1 and Cross 2. The SNP markers were ordered according to their linkage map positions. The correspondences between genotype and phenotype were thoroughly studied.

### BLASTing of SNPs from finemapping

To determine candidate genes, specific KASP marker sequences which are found within the fine mapped interval were BLASTed against the genomes of *M. t.* (LIS - Legume Information System) and of *C. a.* to define their physical positions.

### Analysis of synteny between *Vicia faba* and *Cicer arietinum*

To determine the synteny between *V.f.* and *C.a.* in the interval-of-interest, the syntenic positions of the markers in the final linkage map fragments on the *C. a.* genome were established. For this, the contig DNA sequences from which the new markers were derived and the flanking DNA sequences of the markers from the initial marker set (Webb et al. 2016) were BLASTed against the proteins of the reference sequence of *C. arietinum* (Varshney et al. 2013; NCBI: RefSeq assembly accession no. GCF_000331145.1). BLAST analyses were conducted using the blastx application from the NCBI BLAST+ package (v. 2.7.1). The positions of the transcripts encoding the respective proteins on the reference sequence were extracted from the annotation gff3 file of the sequence in R and assigned to the markers as syntenic positions. To extract all transcripts and the corresponding genes and pro-teins encoded in the region of the *C. a.* genome corresponding to the core region from the annotation file of the reference sequence, the intersect function of bedtools (v2.27.1) was used.

## Results

### SNP marker selection

For the marker-assisted selection, mapping and fine-mapping, markers within the region around the putative VC locus (Khazaei et al. 2015) were chosen and tested. The markers Vf_Mt2g005900_001 and Vf_Mt2g015010_001 from Khazaei et al. (2015) were used to serve as <u>initial boundary markers</u> of the chromosomal to define the initial <u>interval-of-interest</u> and encompass all other chosen markers. In total, six markers from Khazaei et al. (2015), including the initial boundary markers, were chosen. Ten markers from Webb *et al.* 2016 were added since they were determined to be in the interval-of-interest according to the physical placement of the initial boundary markers in *M. t.* genome and the genetic map of Webb *et al.* 2016. Additionally, the marker SNP384 from Song (2017) was included. These 16 markers make up our <u>initial marker set</u> (Table 2) which was used for our first marker assisted selection in the backcross and the construction of <u>draft maps</u> for Cross 1 and Cross 2.

**Table 2.** Seventeen SNP markers from Webb et al. 2016, Song 2017 and D. O’Sullivan (initial marker set). Their function in Cross 1 and 2 and for the backcross is noted. Markers in yellow are the initial boundary markers and encompass the chromosomal interval-of-interest. Blue markers denote the region we suspected the VC gene to be located in. Markers are sorted after Webb et al. (2016).

| SNP-Marker | Usability in families |  |  |
| --- | --- | --- | --- |
|  | Cross 1 | Cross 2 | Backcross |
| Vf_Mt2g005900_001* | X | - | X |
| Vf_Mt1g083330_001* | X | - | - |
| Vf_Mt2g007220_001* | X | - | - |
| Vf_Mt2g007390_001 | - | X | X |
| Vf_Mt2g008150_001 | - | X | - |
| Vf_Mt2g008180_001 | X | - | - |
| Vf_Mt2g008226_001 | X | - | - |
| Vf_Mt2g009230_001 | - | - | - |
| SNP384 | X | X | X |
| Vf_Mt2g009320_001 | X | # | X |
| Vf_Mt2g010740_001* | X | # | X |
| Vf_Mt2g010880_001* | X | - | - |
| Vf_Mt2g010970_001 | - | # | - |
| Vf_Mt2g011080_001 | - | - | X |
| Vf_Mt2g013690_001 | - | X | - |
| Vf_Mt2g013900_001 | - | - | X |
| Vf_Mt2g015010_001 * | - | - | - |

“\*”: Marker from Khazaei et al. 2015; “-” : not polymorphic; “#”: removed from the analysis due to skewed segregation patterns; “X” used in draft maps and final genetic maps and for marker assisted selection

Only subsets of the 16 markers (initial marker set) were polymorphic in the respective genetic backgrounds (**Table 2**). Most notably, the marker Vf_Mt2g015010_001, which served as initial boundary marker, was monomorphic. The polymorphic marker Vf_Mt2g013690_001 was hence chosen as substitute (see Table 2). So, markers Vf_Mt2g005900_001 and Vf_Mt2g013690_001 were subsequently used to define the <u>search frame</u> (Table 3). All markers polymorphic in Cross 1 and Cross 2 were used for the construction of the draft maps. Only three markers were polymorphic in all genetic backgrounds (Table 2, markers SNP384, Vf_Mt2g009320_001 and Vf_Mt2g010740_001) and their chromosomal vicinity was deemed as the most likely region to contain the VC gene, based on the data of Song (2017) and Khazaei et al. (2017), and was therefore the focus for our search for new SNP markers and the VC gene in the transcription analysis. In addition, the markers polymorphic in the back-cross were used for marker assisted selection in the backcrossing program.

**Table 3.** Comparison of LVC vs. HVC data from transcription analysis of Cross 1 and Cross 2. Given are the numbers of detected SNPs and contigs which were found with different transcription analysis methods and search frames.

| Source of SNP and contigs | Area within markers Vf_Mt2g005900_001 and Vf_Mt2g013690_001 (*M.t.* chr. 2; 390,001-3,620,000 bp) |  | Area within markers SNP384 and Vf_Mt2g010740_001 (*M.t.* chr. 2; 1 851 023-2,500,001 bp) |  | Not mappable to *M.t.* |  |
| --- | --- | --- | --- | --- | --- | --- |
|  | SNPs | Contigs | SNPs | Contigs | SNPs | Contigs |
| RNAseq | 393 | 112 | 117 | 30 | 55 | 21 |
| MACE | 128 | 59 | 40 | 17 | 4 | 4 |
| Both, RNAseq + MACE | 73 | 48 | 22 | 12 | 2 | 2 |
| RNAseq & MACE, non-redundant | 448 | 123 | 135 | 35 | 57 | 23 |

Analysis of RNAseq and MACE data sets revealed, in the intersection of Cross 1 and Cross 2, a total of 448 unique SNPs within 123 contigs (Table 3). These contigs could be assigned to chromosome 2 of *M. truncatula*. Of these, 135 unique SNPs from 35 contigs were found within the new interval-of-interst. A total of 73 SNPs were found with both analyses, RNAseq and MACE.

From this data, new markers were developed (Table 4), preferably from those 448 SNP candidates which mapped to the area and within markers Vf_Mt2g005900_001 and Vf_Mt2g013690_001 (*M.t.* chr. 2; 390,001-3,620,000 bp, see Table 3).

**Table 4.**
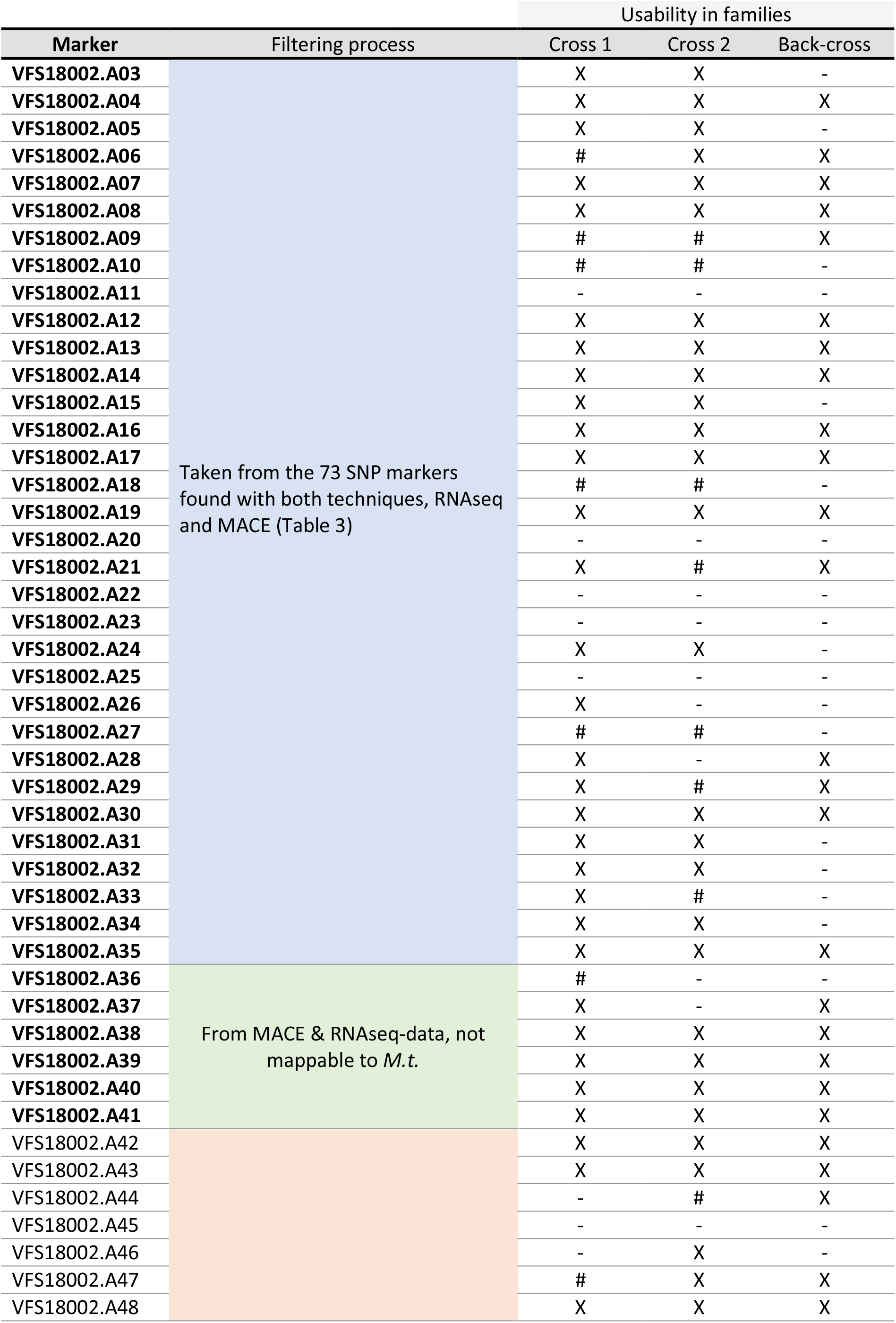
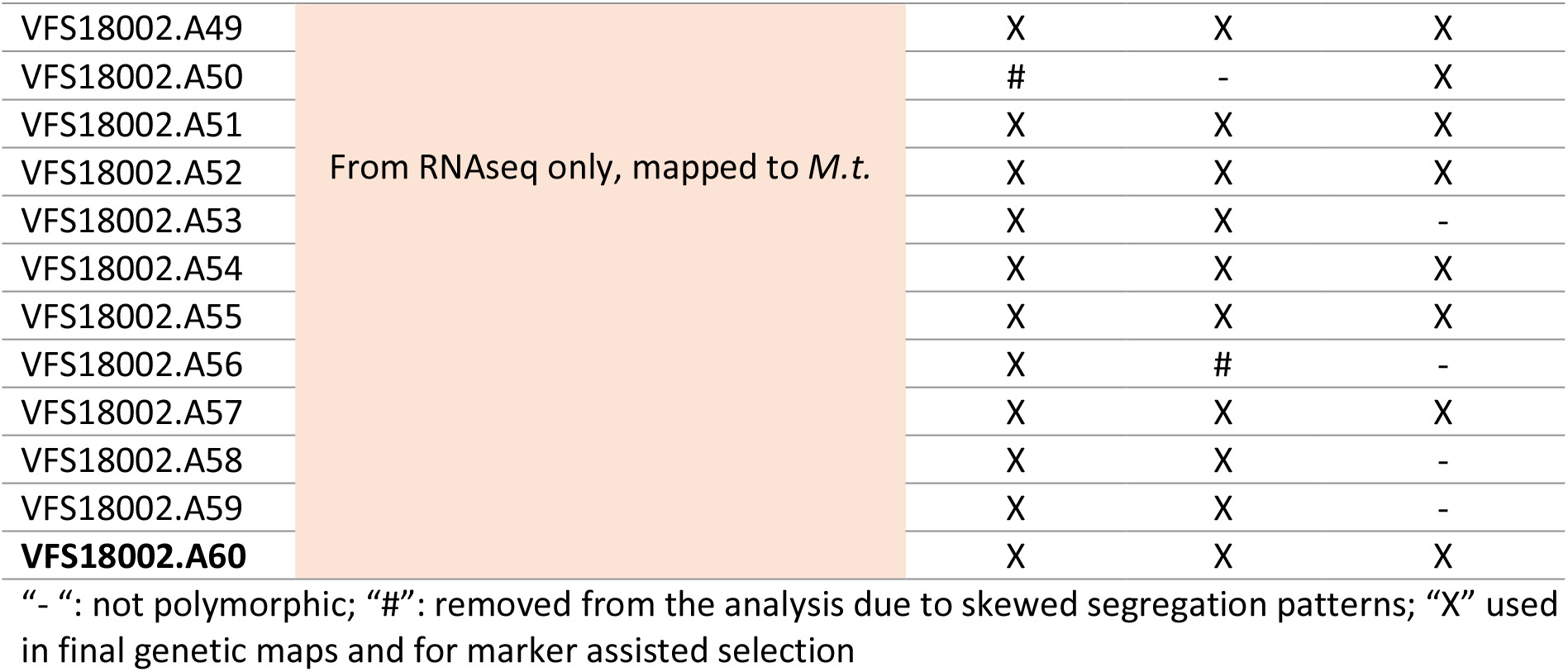
Fifty-eight SNP markers from mRNA analyses RNAseq and MACE. Their origin as well as their function in the three families (Cross 1 and 2 and backcross) is noted.

Of the 58 newly developed markers, 42 markers were used for the construction of the final linkage map fragment and fine-mapping of Cross 1 and 38 markers were used for Cross 2 (see Table 1), since these were polymorphic and did not show skewed segregation patterns. For the backcross, 34 markers were polymorphic and were tested for their suitability for our further winter bean breeding. Those of the initial set markers which were polymorphic for the genetic backgrounds of Cross 1 and 2 plus the above described newly developed markers form the <u>new marker set</u>.

### Linkage maps

For the construction of the draft linkage maps, all F_2_ genotypes present in the F_2_ families were used, i.e., 751 individuals for Cross 1, and 899 individuals for Cross 2. The <u>initial marker set</u> that was employed for draft maps is depicted in Table 2. The marker order in the draft maps followed nearly always the order of their physical placement in the genome sequence of *M. truncatula.* The <u>new marker set</u> gave the final linkage map fragments for Cross 1 and Cross 2.

Only informative F_2_ genotypes were genotyped with the new marker set. Such informative F_2_ genotypes had one or more putative recombination events as determined by their marker genotypes and the draft mapping. A total of 70 individuals for Cross 1 and 74 individuals for Cross 2 were thus reanalyzed with the new marker set. In addition, and as controls, the parental genotypes (10 plants each) of the respective cross were genotyped with the new marker set. The marker data of the remaining F_2_ individuals were inferred based on results from the draft maps.

SNP results of the informative and control individuals were checked for missing data points and for plausibility of their numbers of single and double crossovers. Five F_2_-individuals of Cross 1 and nine of Cross 2 were excluded based on more than 10 missing marker data per individual and on extremely unlikely recombination patterns, such as the apparent occurrence of two or more crossovers in both gametes within 3 cM. This cleansing combined with exclusion of markers with strongly skewed seg-regations (see “SNP marker selection”, Table 4) reduced the length of the map fragments from app. 20 cM in a first, uncleansed approach to between 3 and 6 cM for the final linkage map fragments.

### Fine mapping of the VC locus

For the manual fine-mapping of the sought-for VC-locus, the informative F_2_ plants of Cross 1 and Cross 2 were phenotyped for their VC content.

The resulting VC phenotypes were matched with the genotypes. For this purpose, markers were arranged according to the final linkage map fragments s (Figure 2). The F_2_ individuals were grouped as homozygous HVC, heterozygous and homozygous LVC. Crossovers were thus made visible (see Tables 5A and 5B).

**Figure 2.**
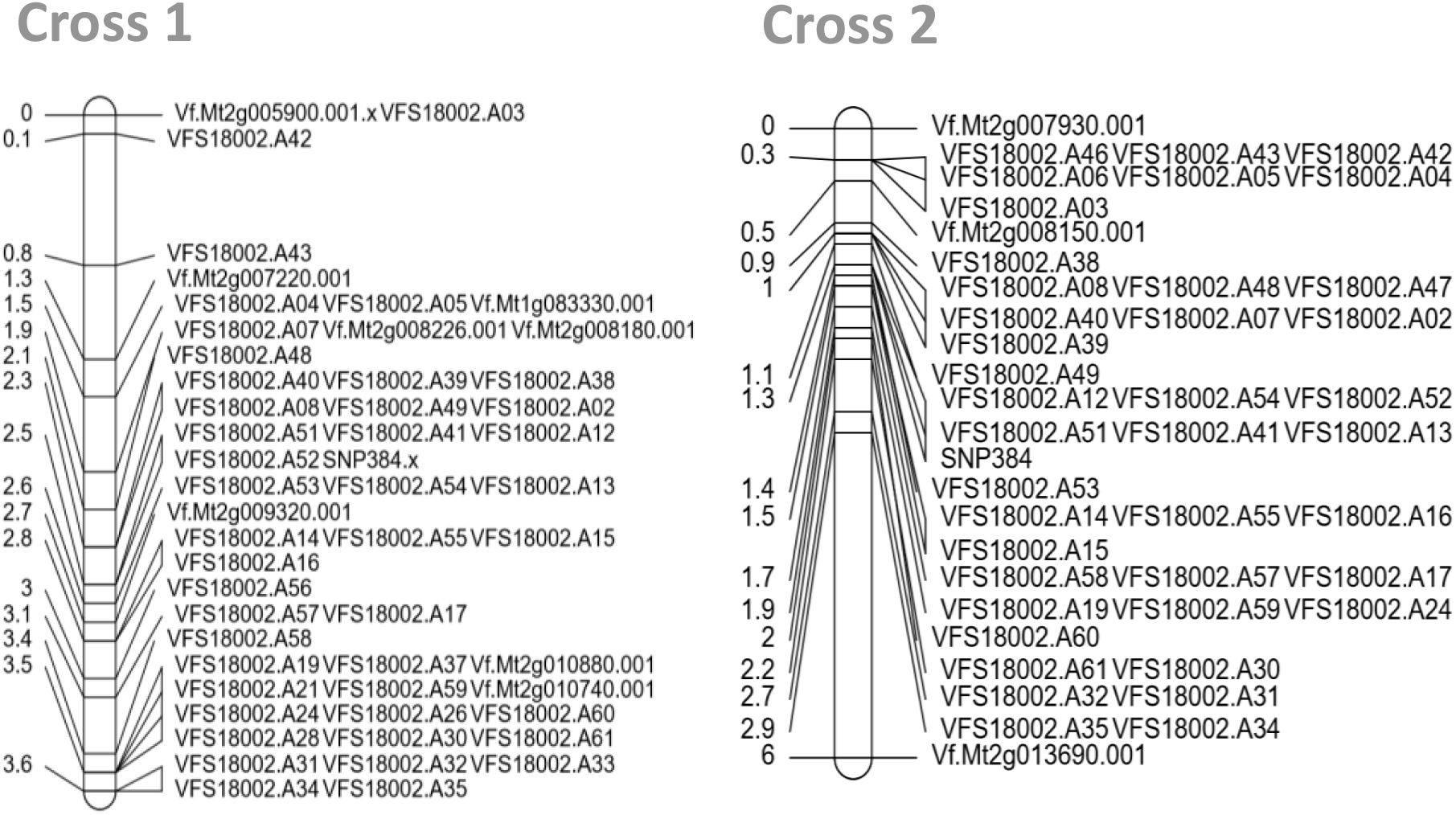
Final linkage map fragments for Cross 1 and Cross 2. Depicted are fragments of chro-mosome 1 of V.f. showing the putative region of the VC locus. Positions of markers are given in cM on the left, marker names for the respective positions are given on the right.

**Table 5A.** Fine-mapping of the vicinity of the VC locus for Cross 1. F_2_ genotypes of the respective individuals are sorted according to their haplotypes, with homozygous for SNP-allele (green, A) as detected in LVC parent on the left to heterozygous (yellow, H) in the middle to homozygous for SNP-alleles (red, B) as detected in HVC parent on the right. The respective HPLC VC contents are given above with the individual numbers of genotypes. Marker names as well as their positions on the genetic linkage map fragments (see Figure 2) are displayed, thus crossovers are visualized. Orange colored marker names define outer boundaries of the interval which, according to this fine-mapping, contains the VC locus. The light green colored marker name depicts the SNP384 which was defined by Khazaei et al. (2017) as a diagnostic marker for VC content.

| Pos. in genetic map (cM) | Marker | F<sub>2</sub> individuales |  |  |  |  |  |
| --- | --- | --- | --- | --- | --- | --- | --- |
|  |  | 587 | 273 | 440 | 765 | 404 | 313 |
|  |  | VC content (HPLC; Total % Vicine/convicine (dry weight)) |  |  |  |  |  |
|  |  | 0.0459 | 0.0436 | 0.4832 | 0.4678 | 0.5824 | 0.5606 |
| Marker genotypes |  |  |  |  |  |  |  |
| 2.28 | VFS18002.A39 | A | H | A | H | B | H |
| 2.28 | VFS18002.A49 | A | H | A | H | B | H |
| 2.48 | VFS18002.A51 | A | A | A | H | B | B |
| 2.55 | VFS18002.A41 | A | A | H | H | B | B |
| 2.55 | VFS18002.A12 | A | A | H | H | B | B |
| 2.55 | VFS18002.A52 | A | A | H | H | B | B |
| 2.55 | SNP384 | A | A | H | H | B | B |
| 2.61 | VFS18002.A54 | A | A | H | H | H | B |
| 2.61 | VFS18002.A13 | A | A | H | H | H | B |
| 2.61 | VFS18002.A53 | A | A | H | H | H | B |
| 2.68 | Vf.Mt2g009320 | A | A | H | H | H | B |
| 2.75 | VFS18002.A14 | A | A | H | A | H | B |
| 2.75 | VFS18002.A55 | A | A | H | A | H | B |

**Table 5B.**
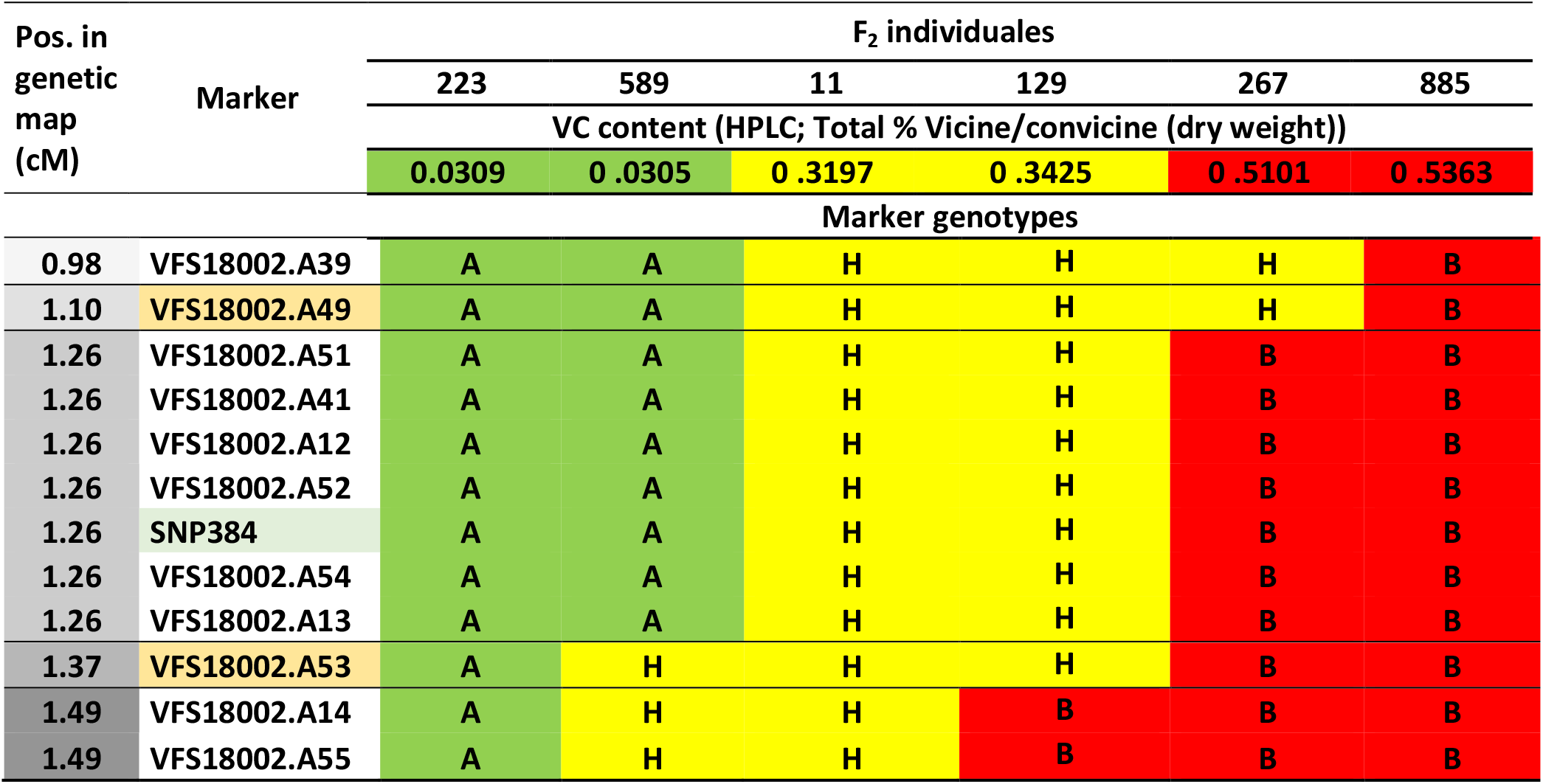
Fine-mapping of the vicinity of the VC locus for Cross 2.

| Pos. in genetic map (cM) | Marker | F<sub>2</sub> individuales |  |  |  |  |  |
| --- | --- | --- | --- | --- | --- | --- | --- |
|  |  | 223 | 589 | 11 | 129 | 267 | 885 |
|  |  | VC content (HPLC; Total % Vicine/convicine (dry weight)) |  |  |  |  |  |
|  |  | 0.0309 | 0.0305 | 0.3197 | 0.3425 | 0.5101 | 0.5363 |
| Marker genotypes |  |  |  |  |  |  |  |
| 0.98 | VFS18002.A39 | A | A | H | H | H | B |
| 1.10 | VFS18002.A49 | A | A | H | H | H | B |
| 1.26 | VFS18002.A51 | A | A | H | H | B | B |
| 1.26 | VFS18002.A41 | A | A | H | H | B | B |
| 1.26 | VFS18002.A12 | A | A | H | H | B | B |
| 1.26 | VFS18002.A52 | A | A | H | H | B | B |
| 1.26 | SNP384 | A | A | H | H | B | B |
| 1.26 | VFS18002.A54 | A | A | H | H | B | B |
| 1.26 | VFS18002.A13 | A | A | H | H | B | B |
| 1.37 | VFS18002.A53 | A | H | H | H | B | B |
| 1.49 | VFS18002.A14 | A | H | H | B | B | B |
| 1.49 | VFS18002.A55 | A | H | H | B | B | B |

It was hence determined which crossovers were closest to the VC locus (see Tables 5A and 5B). The smallest possible SNP interval bearing the sought-for VC locus was determined for both crosses. This interval is referred to as the core region (depicted by orange background of marker names in **Tables 5A and 5B**).

The region between markers VFS18002.A51 and VFS18002.A54 (Cross 1) was 0.14cM smaller than the region between markers VFS18002.A49 and VFS18002.A53 (Cross 2; Table 5A, 5B). All markers in this Cross 1 region were included in the region of Cross 2, and in same order. VFS18002.A51 and VFS18002.A54 of Cross 1 (0.13cM) are consequently the <u>final boundary markers</u> which define the core region where the VC locus must be located.

### SNP marker validation for breeding novel, winterhardy, vc- faba bean lines

Those nine markers which are denominated by the outer boundaries of the intervals in Cross 1 and Cross 2 (see Tables 5A and 5B) were tested in our backcross material to confirm markers which might be good candidates for marker assisted selection. The usability of these markers for the backcross material and the accuracy of the respective marker prediction for the VC content was verified with BC_3_F_2_ genotypes and their respective HPLC results.

Eight of these promising nine markers (see Tables 5A and 5B) proved to be precise selection tools for our backcrossing program, namely markers VFS18002.A49 to VFS18002.A13 (see Table 6) while one marker was not polymorphic here (VFS18002.A53). Two markers did not perfectly match; they were indeed not included in this group of promising markers (markers VFS18002.A14 and VFS18002.A55; see Table 6).

**Table 6.** Markers tested for marker-assisted selection potential in Hiverna/2 * Mélodie/2 backcross material. Markers were selected from the fine-mapping of Cross 1 and Cross 2. They are ordered according to the fine-map of Cross 1.

| Marker | F<sub>2</sub> individuelles of Hiverna/2 * Mélodie/2 backcross generation BC<sub>3</sub> |  |  |  |  |  |  |  |  |  |  |  |
| --- | --- | --- | --- | --- | --- | --- | --- | --- | --- | --- | --- | --- |
|  | VC content (HPLC; Total % Vicine/convicine (dry weight)) |  |  |  |  |  |  |  |  |  |  |  |
|  | 0.0306 | 0.0373 | 0.0396 | 0.0353 | 0.4060 | 0.4079 | 0.4146 | 0.5482 | 0.5539 | 0.5955 | 0.6031 | 0.6285 |
|  | Marker genotypes |  |  |  |  |  |  |  |  |  |  |  |
| VFS18002.A49 | A | A | A | A | H | H | H | B | B | B | B | B |
| VFS18002.A51 | A | A | A | A | H | H | H | B | B | B | B | B |
| VFS18002.A41 | A | A | A | A | H | H | H | B | B | B | B | B |
| VFS18002.A12 | A | A | A | A | H | H | H | B | B | B | B | B |
| VFS18002.A52 | A | A | A | A | H | H | H | B | B | B | B | B |
| SNP384 | A | A | A | A | H | H | H | B | B | B | B | B |
| VFS18002.A54 | A | A | A | A | H | H | H | B | B | B | B | B |
| VFS18002.A13 | A | A | A | A | H | H | H | B | B | B | B | B |
| VFS18002.A53 | Monomorphic in the backcross material |  |  |  |  |  |  |  |  |  |  |  |
| VFS18002.A14 | A | A | A | H | H | H | H | B | B | B | B | B |
| VFS18002.A55 | A | A | A | H | H | H | H | B | B | B | B | B |

### Putative candidate genes from synteny studies and from transcriptome analyses

The core region which was identified via fine-mapping of Cross 1 and Cross 2 was inspected for possible candidate genes. The sequences of the markers covering the core region were BLASTed against the genomes of *M.t.* and *C.a.* (see Table 7) and genes within the core region were noted (see Table 8).

**Table 7.** Markers of core-region and their positions in the *M.t.* genome.

| Position in *V.f.* (cM) | Marker name | Positon in *M.t.* (bp) |
| --- | --- | --- |
| **2.48** | **VFS18002.A51** | 1 764 412 |
| **2.55** | **VFS18002.A41** | no match on Chr. 2 |
| **2.55** | **VFS18002.A12** | 1 828 300 |
| **2.55** | **VFS18002.A52** | 1 852 191 |
| **2.55** | **SNP384** | 1 851 023 |
| **2.61** | **VFS18002.A54** | 1 974 254 |

**Table 8.** Genes located in the region of *M.t.* and *C.a.* syntenic to the core region in *Vicia faba*.

| Gene name | Start position *M.t.* genome (bp) | Product | Assessable in *C.a.* core region equivalent |
| --- | --- | --- | --- |
| MTR_2g009090 | 1 771 882 | Splicing factor 3B subunit, putative | - |
| MTR_2g009110 | 1 780 752 | Splicing factor 3B subunit 1 | Yes |
| MTR_2g009130 | 1 796 938 | Hypothetical protein | - |
| MTR_2g009140 | 1 801 324 | DNA (cytosine-5)-methyltransferase | Yes |
| MTR_2g009150 | 1 807 363 | Transmembrane protein, putative | - |
| MTR_2g009190 | 1 810 285 | Minichromosome maintenance (MCM2/3/5) family protein | Yes |
| MTR_2g009200 | 1 822 351 | Serine carboxypeptidase-like protein | - |
| MTR_2g009220 | 1 828 042 | 2-oxoglutarate/malate translocator | Yes |
| MTR_2g00923 | 1 833 036 | L-ascorbate oxidase | - |
| MTR_2g009270 | 1.848.020 | 3,4-dihydroxy-2-butanone 4-phosphate synthase | Yes |
| MTR_2g009275 | 1 854 017 | Transmembrane protein, putative | Yes |
| MTR_2g009290 | 1 860 925 | Transmembrane protein, putative | - |
| MTR_2g009310 | 1 866 319 | RING-H2 zinc finger protein | Yes |
| MTR_2g009320 | 1 869 086 | Ubiquinol-cytochrome C chaperone protein | Yes |
| MTR_2g009330 | 1 877 274 | Pyruvate decarboxylase | Yes |
| MTR_2g009340 | 1 883 129 | Phototropic-responsive NPH3 family protein | Yes |
| MTR_2g009360 | 1 890 536 | Transcription factor | Yes |
| MTR_2g009390 | 1 899 088 | FKBP-type peptidyl-prolyl cis-trans isomerase | Yes |
| MTR_2g009410 | 1 904 618 | Oligosacccaryltransferase | - |
| MTR_2g009415 | 1 905 205 | tRNA-His | Yes |
| MTR_2g009430 | 1 911 762 | Hypothetical protein | - |
| MTR_2g009450 | 1 915 096 | Leguminosin group485 secreted peptide | Yes |
| MTR_2g009480 | 1 929 386 | Leguminosin group485 secreted peptide | Yes |
| MTR_2g009500 | 1 937 541 | Carbonic anhydrase family protein | Yes |
| MTR_2g009520 | 1 948 713 | Chaperone DnaJ-domain protein | - |
| MTR_2g009530 | 1 952 263 | Hypothetical protein | - |
| MTR_2g009550 | 1 957 898 | ATP synthase subunit beta, putative | - |
| MTR_2g009560 | 1 958 801 | F0F1 ATP synthase subunit beta | - |
| MTR_2g009580 | 1 964 581 | Ulp1 protease family, carboxy-terminal domain protein | Yes |
| MTR_2g009590 | 1 975 139 | Hydroxyproline-rich glycoprotein family protein | - |

A total of 30 genes were thus found in *M.t.*, while a subset of 17 of these could also be found in the corresponding region of *C.a.* (**Table 8**).

The expression analyses revealed that only one gene found in the core-region was differentially expressed (Abo-Vici 2021), namely 3,4-dihydroxy-2-butanone 4-phosphate synthase (or Riboflavin biosynthesis protein, ribBA)).

### Breeding progress of novel, winter hardy, vc- faba bean lines

The backcrossing program, with the selection based on hilum colour and on HPLC results to breed a winter hardy, LVC faba bean line was carried out until 2017 and up to generation BC_3_F_2_. Thereafter, KASP markers were employed (Khazaei et al. 2015; Webb et al. 2015). Starting in 2018, we employed KASP markers derived from our transcription analyses. KASP marker predictions were verified by HPLC results. Our markers allowed to identify VC+/vc− heterozygous BC_4_F_1_ plants and directly employ them for further backcrossing (instead of using BC_4_F_3_), thus we could very markedly speed up the process.

In 2018, the genetic diversity of the backcrossing program was widened. Selected BC_3_F_2_ individuals (Tacke and Link, 2018) were crossed with non-Hiverna/2 winter lines: S_062, S_300, S_306 and S_340 (Gasim and Link, 2007). These lines were chosen for their superior winter hardiness and agronomic performance at Göttingen.

In 2020, we arrived at generation BC_4_F_4_. From this material, four LVC components (see **Table 9**) were sown in October 2019 as generation Syn 0 and in October 2020 as Syn 1, to initiate a novel LVC synthetic variety of winter faba beans as experimental cultivar and as breeding germplasm pool.

**Table 9.** Four LVC components were used to initiate Syn0 of a winter hardy, LVC synthetic population.

| Internal field book numbers | Backcross generation | Detailed pedigree | Component in Syn 0 |
| --- | --- | --- | --- |
| **F19_[622-631]** | BC4F4 | BC4F4(**S_062** * BC3F2(Hiv/2 * Mél/2)) | 1 |
| **F19_[633-638]** | BC4F4 | BC4F4(**S_300** * BC3F2(Hiv/2 * Mél/2)) | 2 |
| **F19_[675-698]** | BC4F4 | BC3F4(**S_306** * BC2F2(Hiv/2 * Mél/2)) | 3 |
| **F19_[702-720]** | BC3F4 | BC3F4(**S_340** * BC2F2(Hiv/2 * Mél/2)) | 4 |

## Discussion

In 2015, Khazaei et al. published a genetic map of the fragment of faba bean chromosome 1 which most likely contains the VC locus. This was the first mapping attempt of this locus and the VC gene. It reduced the search area for the causative gene down to approximately 4cM. For this, the authors employed a set of 210 F_5_ recombinant inbred lines (RILs) of a cross between Mélodie/2, a LVC inbred line, and ILB938/2, a HVC inbred line. Genotyping was done with a set of 188 polymorphic SNPs (Khazaei et al. 2015).

Gutiérrez et al. (2016) reported corresponding findings from their cross Vf6 * 1268. The putative region of the respective locus in their approach amounts also to app. 4cM. We employed two crosses with two different genetic backgrounds containing 751 and 899 individuals, respectively. Sixteen markers from Webb et al. (2016) and Song (2017) allowed the draft mapping. Via mRNA analyses, 58 new markers were developed, 42 and 38 of which were polymorphic for the two genetic backgrounds and were thus used for the creation of final genetic linkage map fragments. The two resulting maps show, for those markers which were also used by Khazaei et al. (2017), the almost same order. The so far empty gaps between those markers are now enriched with 42 and 38 new markers, respectively. The subsequent fine mapping resulted in a core region for the VC gene of app. only 0.13 cM.

The relatively small area of this core region yielded a set of putative candidates. Additionally, our approach towards a candidate gene via expression analysis also yielded a set of candidates, which were differentially expressed in immature seed coat tissue of our LVC and HVC NIL pairs.

The overlap between the two approaches clearly indicated that the most likely VC candidate gene was 3,4-dihydroxy-2-butanone 4-phosphate synthase, or bifunctional riboflavin biosynthesis protein RIBA 1. Recently, Björnsdotter et al. (2020) named the gene RIBA 1 as the putative candidate gene for the vc- phenotype. Their findings strongly suggest that vicine and convicine are side products of the riboflavin biosynthesis from the purine GTP. Their hypothesis for the cause of the LVC phenotype is a frame shift insertion in the RIBA1 enzyme. They accordingly named the gene for the RIBA1 enzyme VC1. Our findings seem to verify the conclusion of Björnsdotter et al. (2020).

Additionally, the mRNA analyses lead to a set of new markers in the VC core region, ready for use for marker assisted selection. We employed them and bred new, LVC winter faba bean lines, aiming at highly versatile feed stuff from highly productive winter beans. Syn 1 of the first, novel LVC winter faba bean variety is currently (2021 season) in the field. This introduction of the LVC feature will make winter faba beans more attractive for farmers and feed producers.

## Acknowledgements

We very thankfully acknowledge funding of this project by BLE/BMEL (Abo-Vici, FKZ 2815EPS004). We acknowledge the very helpful donations of F_5_-individuals of Cross 1 (University Helslinki) and of Cross 2 (NPZ Lembke KG) to the Abo-Vici project. Donal O’Sullivan and Deepti Angra gave us the initial marker set; we thank them indeed. We would like to thank Helen Appleyard at NIAB for the very many HLPC analyses and very efficient cooperation. Finally, we would like to express our thankfulness to the team at Georg-August University of Göttingen, and especially to Regina March and Sonja Yaman.

